# Evolution of age-specific decline in stress phenotypes is driven by both antagonistic pleiotropy and mutation accumulation

**DOI:** 10.1101/115931

**Authors:** Elizabeth R. Everman, Theodore J. Morgan

**Author notes:** Corresponding Author: Theodore J. Morgan, 116 Ackert Hall, Division of Biology, Kansas State University, Manhattan, KS 66506.

## Abstract

Efforts to more fully understand and test evolutionary theories of aging have produced distinct predictions for mutation accumulation (MA) and antagonistic pleiotropy (AP) mechanisms. We build on these predictions through the use of association mapping and investigation of the change in additive effects of polymorphisms across age and among traits for multiple stress response phenotypes. We found that cold stress survival with acclimation, cold stress survival without acclimation, and starvation resistance declined with age and that changes in the genetic architecture of each phenotype were consistent with MA predictions. We used a novel test for MA and AP by calculating the additive effect of polymorphisms across ages and found support for both MA and AP mechanisms in the age-related decline in stress tolerance. These patterns suggest both MA and AP contribute to age-related change in stress response and highlight the utility of association mapping to identify genetic shifts across age.

## Introduction

The intensity of natural selection changes over an organism’s lifespan, having greatest effect early in life, as individuals reach reproductive maturity, and smaller effect as organisms age (Charlesworth, 2001; Charlesworth and Hughes, 1996; Fisher, 1930; Haldane, 1941; Hamilton, 1966; Medawar, 1952; Williams, 1957). Decreased effectiveness of natural selection at old age results in the accumulation of deleterious polymorphisms in populations and leads to decline in age-specific fitness, characteristic of senescence (Hamilton, 1966; Medawar, 1952; Williams, 1957). Senescence is expected to negatively impact phenotypes related to fitness and is thought to have evolved through two non-mutually exclusive genetic mechanisms (Bowler and Terblanche, 2008; Charlesworth, 1994; Ricklefs and Finch, 1995). Under mutation accumulation (MA; Medawar, 1952), decreased effectiveness of natural selection over lifespan allows the retention of deleterious polymorphisms that are only expressed later in life (Ricklefs and Finch, 1995). Under antagonistic pleiotropy (AP; Williams, 1957), genes that are expressed over a wide window of an individual’s lifespan have positive effects on fitness at young age and negative effects on fitness at old age (Charlesworth, 2001; Maklakov et al., 2015; Ricklefs and Finch, 1995; Williams, 1957). Both mechanisms rely on the relaxation of natural selection later in an organism’s life but have unique predictions for how age-dependent genetic control of phenotypes changes.

Age-related declines and the influence of the MA and AP mechanisms have been well documented for life-history phenotypes such as mortality and fecundity (Bowler and Terblanche, 2008; Charlesworth and Hughes, 1996; Durham et al., 2014; Engström et al., 1989; Hughes et al., 2002; Pletcher et al., 1998; Promislow et al., 1996; Rose, 1984; Rose et al., 1992; Snoke and Promislow, 2003; Tatar et al., 1996), but far less is known about how stress response phenotypes change with age (Bowler and Terblanche, 2008). Stress response over an organism’s lifespan is a critical component of fitness, and is an important modulator of lifespan (Colinet et al., 2015). Variation in stress response can influence the persistence and evolution of populations over short time scales, especially in variable environments (Bergland et al., 2014). In species that experience seasonal change in thermal regime, changes in the demographic structure of populations can also drastically influence the ability of individuals to tolerate stressful temperatures. For example, in *Drosophila melanogaster* and *D. simulans,* populations are primarily composed of young individuals in the spring when temperatures are increasing on average and primarily of older individuals in the fall when temperatures are decreasing on average (Behrman et al., 2015). Thus, measures of thermal tolerance at one point in the season therefore do not reflect the influence of seasonal variation in age on thermal tolerance. Such shifts in the age structure of populations coupled with age related changes in the genetic control of fitness phenotypes have the potential to dramatically influence short- and long-term responses to environmental variation.

The MA and AP aging mechanisms make predictions about age-specific changes in multiple quantitative genetic parameters. Under MA, genetic variance is expected to increase with age because of the expression of age-restricted polymorphisms (Charlesworth, 2001; Charlesworth and Hughes, 1996; Hughes et al., 2002; Leips et al., 2006). These late acting polymorphisms are retained in the population because the individuals that possess them have successfully reproduced, allowing such alleles to evade natural selection (Charlesworth, 2001; Haldane, 1941; Maklakov et al., 2015). Additionally, because the genetic control of the phenotype across ages is independent, the genetic correlation of the phenotype between young and old individuals is expected to be non-negative (Charlesworth, 2001; Maklakov et al., 2015; Reynolds et al., 2007). In contrast, under the AP hypothesis, age-related change in phenotypes is the result of a genetic trade-off, where polymorphisms that are beneficial early in life are detrimental late in life (Leips et al., 2006; Maklakov et al., 2015). These polymorphisms are retained in the population because of their beneficial effects on fitness at young age (Charlesworth and Hughes, 1996; Leips et al., 2006; Maklakov et al., 2015). AP does not make clear predictions for changes in variance components with age; however, because the same polymorphisms are expected to influence the phenotype with opposite effects across age, the genetic correlation across ages is expected to be negative (Charlesworth and Hughes, 1996; Hughes et al., 2002; Leips et al., 2006).

Despite these clear predictions, the influence of MA and AP and the theory behind these mechanisms does not fully explain age-related change in phenotypes. One reason for this is that the signature of AP may be lost because of small effects or fixation of loci involved in AP leading to ascertainment bias toward MA (Maklakov et al., 2015; Moorad and Promislow, 2009). Further, calculations of genetic variance and correlations are indirect metrics of age-related change in the genetic control of phenotypes (Charlesworth, 2001). In contrast, the use of association mapping allows us to extend these predictions to more explicitly evaluate subtle patterns predicted by the MA and AP theories of aging. Durham et al. (2014) previously used association mapping to identify and compare polymorphisms that are associated with variation in phenotypes measured at multiple ages. This use of association mapping can be extended in two important ways. First, sets of associated polymorphisms can be compared between two different phenotypes across age as well as within phenotypes across age. Second, even if sets of significantly associated polymorphisms are non-overlapping, association mapping allows the evaluation of age-related shifts in the additive effects of associated polymorphisms, thus facilitating the detection of weak antagonistic effects across age or phenotypes.

As an example, consider a hypothetical phenotype measured in young and old individuals that is associated with non-overlapping sets of polymorphisms at each age. Two different polymorphisms are associated with the hypothetical phenotype at young age and have positive additive effects on the young phenotype. The polymorphism that is consistent with MA will shift from a significant positive additive effect at young age to an effect that is near zero or of the same sign at old age. In contrast, the polymorphism that is consistent with AP will shift from a positive additive effect at young age to a negative additive effect at old age. Thus, even though association mapping may not detect the antagonistic polymorphism with small effect in old individuals, calculation of additive effects of polymorphisms across age can be used to detect signals of AP (Fig. 1; Maklakov et al., 2015).

**Figure 1.**
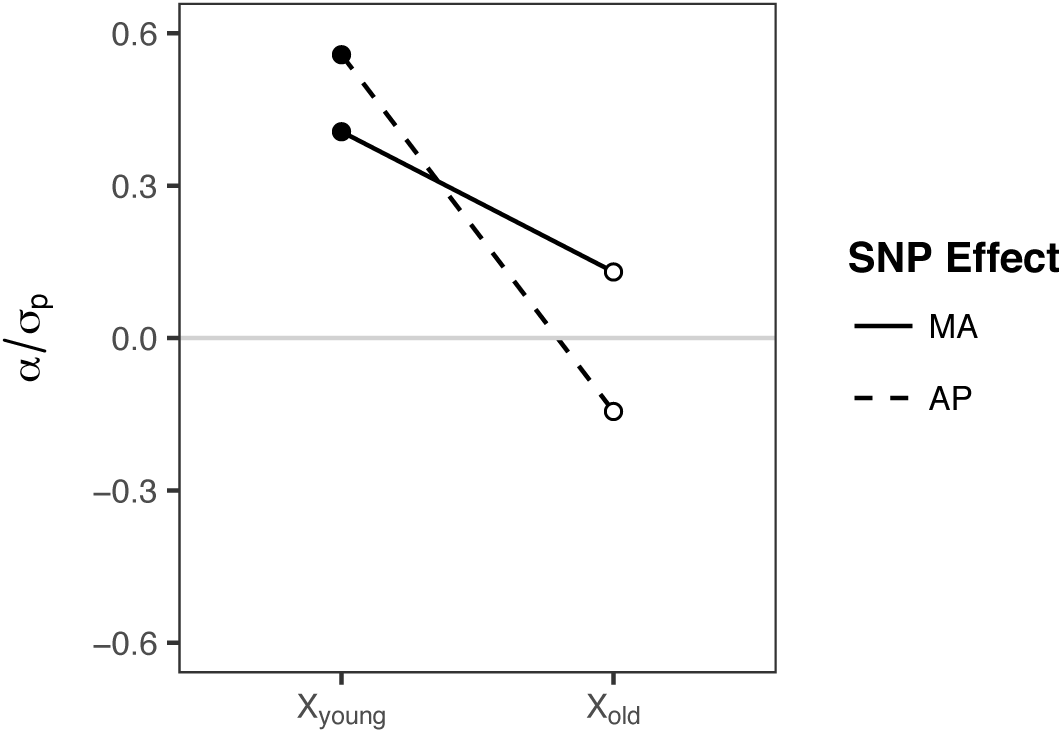
An example of how calculated additive effects (α/σ_p_) of polymorphisms identified through association mapping can be used to characterize the role of mutation accumulation (MA) and antagonistic pleiotropy (AP) on the age-related change in hypothetical phenotype *X.* Two polymorphisms that are associated with phenotype *X* in young individuals (*X_young_*) but not in old individuals (*X_old_*). The additive effects of the associated polymorphisms at young ages are represented by the closed symbols. The open symbols indicate the additive effects of *X_young_* polymorphisms on the phenotype at old age (*X_old_*). The polymorphism connected with a solid line demonstrates the shifting effect across age consistent with MA, where the polymorphism has a small additive effect of the same sign (positive) on the phenotype at old age (*X_old_*) as the effect of the polymorphism on *X_young_.* The polymorphism connected with a dashed line illustrates an AP pattern, where the polymorphism has an additive effect that is of the opposite sign (negative) on *X_old_* compared to the effect of the polymorphism on *X_young_*

In the current study, we used a combination of association mapping and quantitative genetic analysis to dissect the variation in age-related changes in four environmental stress response phenotypes and tested the influence of MA and AP. To do this, we measured age related survival after cold-stress with acclimation and without acclimation, thermal phenotypic plasticity, and starvation resistance in a genetically diverse *D. melanogaster* mapping population from the *Drosophila* Genetic Reference Panel (DGRP; Huang et al., 2014; Mackay et al., 2012). Starvation resistance and cold tolerance are quantitative genetic phenotypes, with heritable variation existing for each (Bowler and Terblanche, 2008; Gerken et al., 2015; Hoffmann et al., 2005; Imasheva et al., 1998; Mackay et al., 2012; Morgan and Mackay, 2006; Overgaard et al., 2010; Schwasinger-Schmidt et al., 2012). However, much of the previous work examining age related change in cold tolerance has been done in a single genotype (Czajka and Lee, 1990), and little research is available to inform how starvation resistance will change with age (but see (Colinet et al., 2015). We predicted that starvation resistance and cold tolerance (measured as acclimation and non-acclimation survivorship) would decline with age.

Genetic variation has also been documented for various forms of thermal phenotypic plasticity (Fallis et al., 2014; Gerken et al., 2015). Short-term acclimation through rapid cold-hardening (RCH score; Lee et al., 1987) is one form of plasticity that occurs when organisms are exposed to a mild thermal stress before experiencing more stressful conditions (Coulson and Bale, 1990; Czajka and Lee, 1990; Gerken et al., 2015; Lee et al., 1987; Powell and Bale, 2005). In flies and other ectothermic species, this pre-treatment usually results in increased cold survivorship and provides a simple model of the physiological response of ectotherms as they respond to episodic fluctuations in temperature (Bozinovic et al., 2011; Coulson and Bale, 1990; Huey et al., 2012; Ju et al., 2011; Niehaus et al., 2012). In most cases, a strong, beneficial acclimation response is detected in young adult individuals (Gerken et al., 2015; Ju et al., 2011; Powell and Bale, 2005; Rajamohan and Sinclair, 2009), but age-related changes in this form of phenotypic plasticity have not been examined. We expected flies to lose the ability to survive acclimation and non-acclimation cold stress at a similar rate, resulting in acclimation scores (the difference in survival with and without cold acclimation) that would not change with age.

## Methods

### Fly stocks

The *Drosophila* Genetic Reference Panel (DGRP) was established as a set of natural isogeneic lines founded from a single population in Raleigh, NC (Table S1; Mackay et al., 2012). Stocks were obtained from Bloomington Stock Center and maintained at 25°C on a 12-hour light-dark cycle on cornmeal-molasses agar sprinkled with active yeast. Parents of experimental flies were sorted over light CO_2_ anesthesia and placed into vials containing five individuals of each sex to establish the first experimental block. Females were allowed to mate and lay eggs for three days, after which the parents were transferred to a new set of vials. Egg laying continued in the new vials for three days to establish the second experimental block, and then parents were discarded. Experimental flies were collected on the third day of eclosion and sorted by sex to a density of 10 same-sex individuals per vial. Our experimental design measured responses in “young” and “old” cohorts of flies. Young flies were aged for 1 week (7 days) at 25°C, while old flies were aged for four weeks (28 days) at 25°C. The “old” cohort timing was selected because this was an advanced age time point, but prior to a significant decline in average survivorship among lines (Ivanov et al., 2015). Previous research has also demonstrated reduced fecundity at this age (Leips et al., 2006; Tatar et al., 1996). Experimental flies in the “old” cohort were tipped every third day to new media until flies were tested at 28 days.

## Age-related stress responses

### Cold stress responses

We measured three cold stress responses on 101 DGRP lines at young and old age (Table S1). We measured acclimation survival using a rapid cold hardening treatment that consisted of a two-hour exposure to 4°C immediately prior to cold shock at −6°C for one hour (Gerken et al., 2015; Lee et al., 1987). Following cold shock, the flies were transferred to fresh media and allowed to recover at 25°C for 24 hours (Fig. S1A). We also measured non-acclimation survival by transferring flies directly (without acclimation) to −6°C for one hour (Fig. S1B). As with the acclimation treatment, the non-acclimated flies were transferred to fresh media following cold stress and allowed to recover for 24 hours at 25°C. After 24 hours, the proportion of flies that had survived each treatment was recorded by counting the number of individuals in each vial that were capable of coordinated movement (flying or walking). Acclimation score (or rapid cold hardening capacity) was calculated by subtracting non-acclimation survivorship from acclimation survivorship (Gerken et al., 2015). A total of four replicates per sex, line, age, and cold stress treatment were measured in two experimental blocks with 10 individuals per vial replicate.

### Starvation resistance

We measured starvation resistance in 164 DGRP lines, including the 101 lines used in the cold stress response experiments, at young and old age (Table S1). Young and old flies were maintained on standard media until they were one or four weeks of age, respectively. At one or four weeks of age, flies were transferred to starvation media (1.5% agarose) and maintained at 25°C. Vials were monitored every four hours, and average time of death per vial was recorded as the response. A total of three replicates per sex, line, and age were measured with 10 individuals per vial replicate.

### Cold-stress responses in physiologically aged individuals

We conducted an additional experiment to test for variability in the rate of senescence among DGRP lines. For this experiment, 10 lines were randomly selected from the 101 included in the cold stress experiment (Table S1), and experimental flies were obtained as described previously. Physiologically-aged flies were set up at a density of 20 individuals per vial and maintained at 25°C until the number of flies per vial and line reached approximately 50% (Td50). At this point, acclimation survivorship, non-acclimation survivorship, and acclimation score were measured for the physiologically aged flies and compared to that of the chronologically aged (four-week-old) flies.

## Data analysis

### Genetic variation

Genetic variation among all lines was analyzed via mixed-model ANOVA for the cold stress responses (acclimation survivorship, non-acclimation survivorship, and acclimation score) and starvation resistance. The model for each analysis included the main effects of age and sex, as well as interactions, with block (for cold stress responses only) and line as random effects. Specific effects of sex by age interactions were tested with Tukey’s HSD post hoc test, with an experiment-wide α = 0.05.

To assess the effect of variation in rate of aging on cold stress tolerance (i.e. physiological vs chronological cold stress phenotypes), linear regression was used to compare the mean cold stress responses of four-week-old flies with cold stress response of flies at their line specific Td50. However, because none of the physiologically-aged flies survived non acclimated cold stress, we only compared acclimation survivorship in this analysis.

### Genome-wide association analysis

We used association mapping to identify regions of the genome that were significantly associated with variation in acclimation survivorship, non-acclimation survivorship, acclimation score, and starvation resistance. Association mapping was performed on each age and phenotype separately, and significance was assigned at −log_10_(5) (Durham et al., 2014; Gerken et al., 2015; Mackay et al., 2012). Shifts in genetic architecture across age and phenotype were assessed by comparing the significant polymorphisms associated with each age-specific phenotype. We performed gene ontology (GO) enrichment analysis using FlyMine (Lyne et al., 2007) to determine whether specific classes of genes or pathways were overrepresented in the loci associated with each phenotype and age.

### Quantitative genetic analyses

Heritability, variance components, and genetic correlations were estimated using the program H2boot, which applies bootstrap resampling to quantitative genetic data (Phillips, 1998). Acclimation survivorship, non-acclimation survivorship, acclimation score, and starvation resistance for one- and four-week-old flies were treated as eight phenotypes. Data were analyzed using a one-way ANOVA, resampling lines 10,000 times with replacement. Because DGRP lines are inbred homozygous lines, reported heritability estimates are broad sense, and were estimated as:

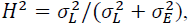

where 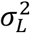 is the among line homozygous genetic variance component, and 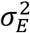 is the environmental variance component. The coefficient of homozygous genetic variance was used to assess the effect of age on changes in homozygous genetic variance, and was estimated as:

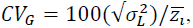

where z¯_*l*_ is the phenotype mean. Genetic correlations across ages were estimated as:

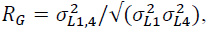

where 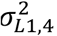 is the covariance component of the phenotype across the two ages tested, 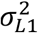 is the among line homozygous genetic variance component of the phenotype for one-week-old flies, and 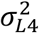 is the among line homozygous genetic variance component of the phenotype for four week-old flies. Variance component and genetic correlation estimates were reported as the average of the 10,000 bootstraps, and the variation in the estimate was used to generate standard errors for each term.

The additive effect of an allele (α) for associated polymorphisms was calculated as one half the difference in phenotypic mean of lines grouped according to homozygous genotype, corrected for *Wolbachia* infection and TE insertions (Falconer and Mackay, 1996; Huang et al., 2014):

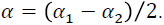

Allele class was designated as major if allele frequency exceeded 50% in the experimental population. Standardized allele effects were calculated as the additive effect divided by the standard deviation of the phenotype:

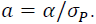

In the DGRP, evidence of MA and AP was examined four ways. First, if MA contributes to population-level decline in a phenotype, late acting alleles will inflate *CV_G_* in four-week-old flies. Second, if MA is responsible for age-specific decline in phenotypes, unique regions of the genome should be associated with the phenotype at young and old age. Under AP, regions of the genome that are associated with the phenotype in young and old individuals should overlap. Third, under MA, we expect the genetic correlation between ages for each phenotype to be non negative, due to the expectation that the additive effects at different ages are independent (Hughes et al., 2002; Leips et al., 2006), while under AP, we expect the genetic correlation between ages for each phenotype to be negative because regions of the genome associated with the phenotype in young and old individuals overlap (Charlesworth and Hughes, 1996; Maklakov et al., 2015). Finally, under MA, the additive effects of polymorphisms that are associated with a phenotype in young individuals are expected to have additive effects on the phenotype in old individuals that are smaller but of the same sign (Fig. 1), while under AP, polymorphisms associated with a phenotype in young individuals are expected to have additive effects on the phenotype in old individuals that are of the opposite sign (Fig. 1).

In addition to testing these predictions of MA and AP within each phenotype across age, we also tested the role of MA and AP in age-related change between phenotypes. As above for each phenotype, we assessed the level of overlap of polymorphisms and genetic correlations between phenotypes and calculated the additive effects of polymorphisms associated with each phenotype on every other phenotype and age in our study. For example, the additive effects of polymorphisms associated with acclimation survival at one week were calculated for both the one and four-week non-acclimation survival and starvation resistance responses.

### Tests for selection

We used the QTL sign test (QTLST) to the test the direction of the additive effects of associated polymorphisms identified through association mapping. The QTLST was developed by (Orr, 1998) to determine whether the signs of QTL effects were indicative of directional selection acting on a phenotype. The probability for rejecting the null hypothesis that selection does not influence the phenotype was calculated as in Orr (1998):

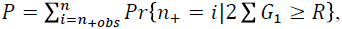

In this study, we treated n as the number of associated polymorphisms detected for each phenotype, n_+obs_ as the number of these polymorphisms that had a positive additive effect, and G as a vector of all additive effects. Because association mapping does not involve generation of a mapping population from distinct lines, R was simply the standard deviation of the phenotype of the population. An exponential distribution of the polymorphism effects was assumed in our adaptation of this model as in applications of QTLST to QTL data. Because QTLST is sensitive to high variance among additive effects, we also performed the QTLST-EE, which assumes that each of the additive effects are equal (Anderson and Slatkin, 2003; Rice and Townsend, 2012). Analyses were done in R using code adapted from Muir et al. (2014) (Hope, 2013; Muir et al., 2014; R Core Team, 2015; Wickham, 2009).

## Results

### Phenotypic responses

All phenotypes measured in this study were variable across ages, lines, and sexes (Fig. 2; Table S2). Two-way interactions among these effects were also significant, except for age by sex in non-acclimation survivorship (Table S2). The three-way interaction between age, line, and sex explained a significant amount of variation for acclimation survivorship and starvation resistance as well (Table S2). On average, acclimation and non-acclimation survivorship and starvation resistance decreased with age as expected (Fig. 2A, D, J; Table S2). However, age-related decline was stronger in non-acclimation survivorship compared to acclimation survivorship, resulting in an average acclimation score that increased significantly with age (Fig. 2G; Table S2). When the sex-specific cold tolerances were analyzed, male flies maintained their capacity to survive the acclimation treatment across age (Fig. 2B; adj. *P* = 0.64). However, this maintenance across age was not observed in the non-acclimation treatment (Fig. 1E; adj. *P* < 0.001). In females, flies tended to lose the survival capacity at an equal rate for both acclimation and non acclimation treatments (Fig. 2B, E). As a result of these sex-specific age-related responses, female acclimation score did not change with age (adj. *P* = 0.46; Fig. 2H), but male acclimation score increased (adj. *P* < 0.001). Thus, the population-level increase in acclimation score was likely driven by retention of cold tolerance in the acclimated male flies. Post hoc comparisons of sex-specific starvation resistance revealed that the age-related average decrease in starvation resistance was primarily driven by a significant decrease in resistance in females (adj. p < 0.001; Fig. 2K).

**Figure 2.**
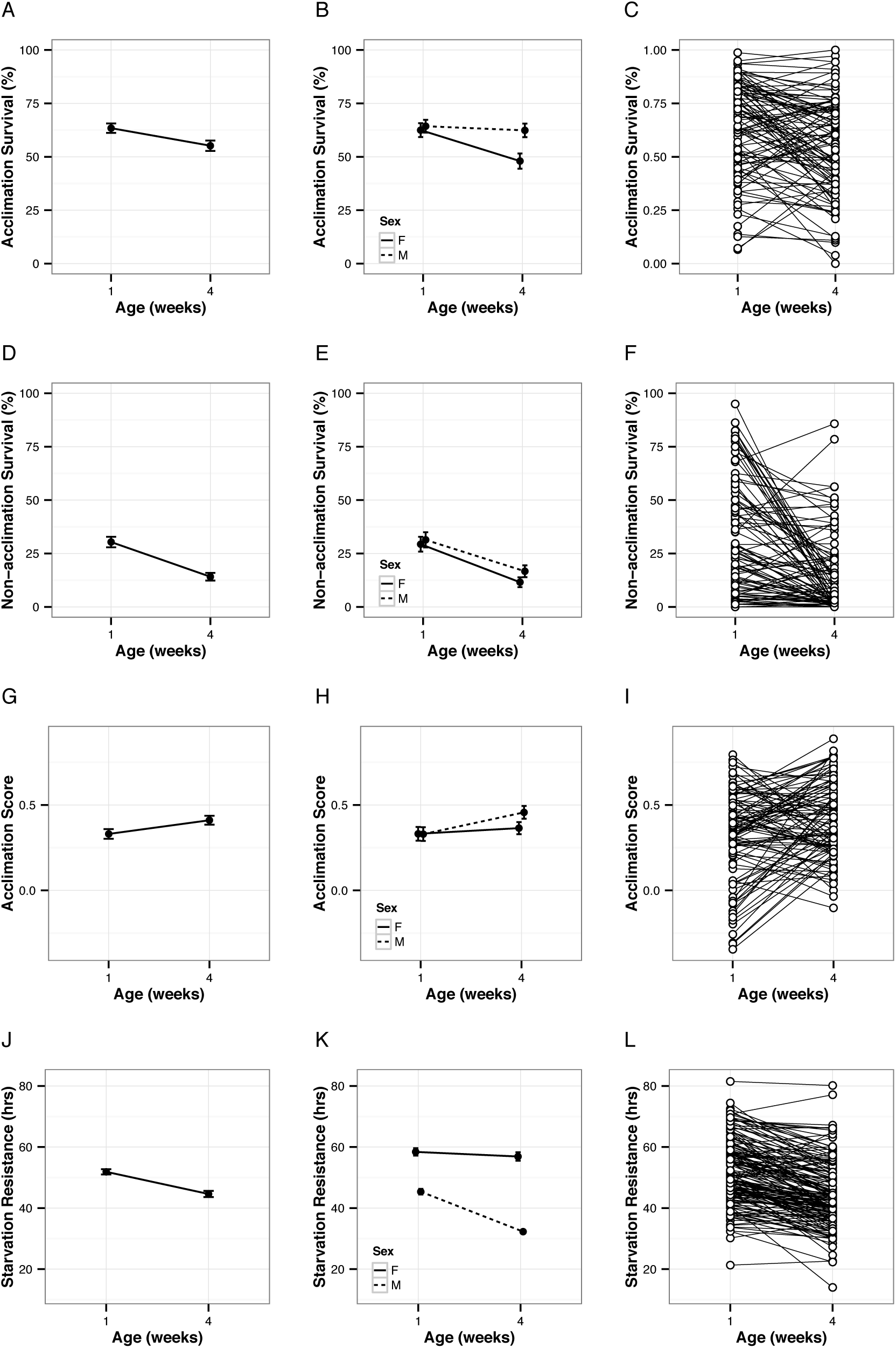
Plots of mean phenotypes across age are shown with 95% CI for the total population (left column) and by sex (middle columns); among-line variation observed for each phenotype is shown as line averages (right column). A. Average acclimation survival declined significantly with age (*F_1,1212_ =* 49.5, *P* < 0.001). B. Males retained their ability to survive the acclimation treatment, while acclimation survival significantly declined in females (age by sex: *F_1,1212_ =* 28.9, *P* < 0.001). C. Acclimation survival significantly varied among the 101 DGRP lines (age by line: *F_100,1212_ =* 3.21, *P* < 0.001). D. Average non-acclimation survival declined significantly with age (*F_1,1212_ =* 215.4, *P* < 0.001). E. Non-acclimation survival significantly declined in both females and males to a similar degree (age by sex: *F_1,1212_ =* 1.98, *P* = 0.16). F. Non-acclimation survival significantly varied among the 101 DGRP lines (age by line: *F_100,1212_ =* 5.88, *P* < 0.001). G. Average acclimation score increased significantly with age (*F_1,1212_ =* 25.13, *P* < 0.001). H. Acclimation score significantly increased in males, but remained consistent across age for females (age by sex: *F_1,1212_ =* 8.68, *P* < 0.01). I. Acclimation score significantly varied among the 101 DGRP lines (age by line: *F_100,1212_ =* 3.18, *P* < 0.001). J. Average starvation resistance decreased significantly with age (*F_1,1623_ =* 893.0, *P* < 0.001). K. Starvation resistance significantly decreased in both sexes, but to a larger degree in females (age by sex: *F_1,1623_ =* 567.0, *P* < 0.001). L. Starvation resistance significantly varied among the 164 DGRP lines (age by line: *F_163,1623_ =* 6.0, *P* < 0.001).

Genotype-specific responses for each phenotype were highly variable (Fig. 2, right column; Table S3); an age-related increase in stress resistance was observed for some lines, while responses in other lines remained constant or decreased (Fig. 2, right column; Table S2). Negative acclimation scores were obtained for several lines screened at one week of age, suggesting that the acclimation treatment had a detrimental effect on survivorship; however, the vast majority of lines responded positively to this treatment (Fig. 2I). When screened at four weeks, the negative acclimation effect largely disappeared as only two lines had acclimation scores below 0. This change in the pattern of cold tolerance with age may have important implications for the role of plasticity in maintaining stress response with age.

### Variation in senescence

To assess the relationship between chronological and physiological age, we measured acclimation survival, non-acclimation survival, and acclimation score on ten randomly selected lines from the DGRP (Table S1). Non-acclimation survival of physiologically-aged flies was 0 in all lines, so only acclimation survivorship was compared between the physiologically-aged (Td50) flies and the chronologically-aged (four-week-old) flies. The average acclimation survivorship of the ten lines selected for the physiological aging experiment was comparable to that of the 101 lines. The average proportion survived following acclimation for one-week old flies from the ten lines was 74.0 ± 8.1 S.E. compared to 63.4 ± 1.1 S.E. in 101 lines, while the average proportion survived following acclimation in four-week old flies was 52.2 ± 9.9 S.E. compared to 55.2 ± 1.2 S.E. in 101 lines. Regression analysis indicated that acclimation survivorship measured in four-week old flies was a good predictor of acclimation survivorship in flies that reached a similar physiological age (R = 0.83; *P* < 0.003; Fig. 3; Table S4). While longevity does vary among DGRP lines, variation in lifespan did not significantly alter the rank order of acclimation survival among the lines. This suggests that variation in longevity among the DGRP lines does not influence the age-related change in detected in phenotypes between young and old flies.

**Figure 3.**
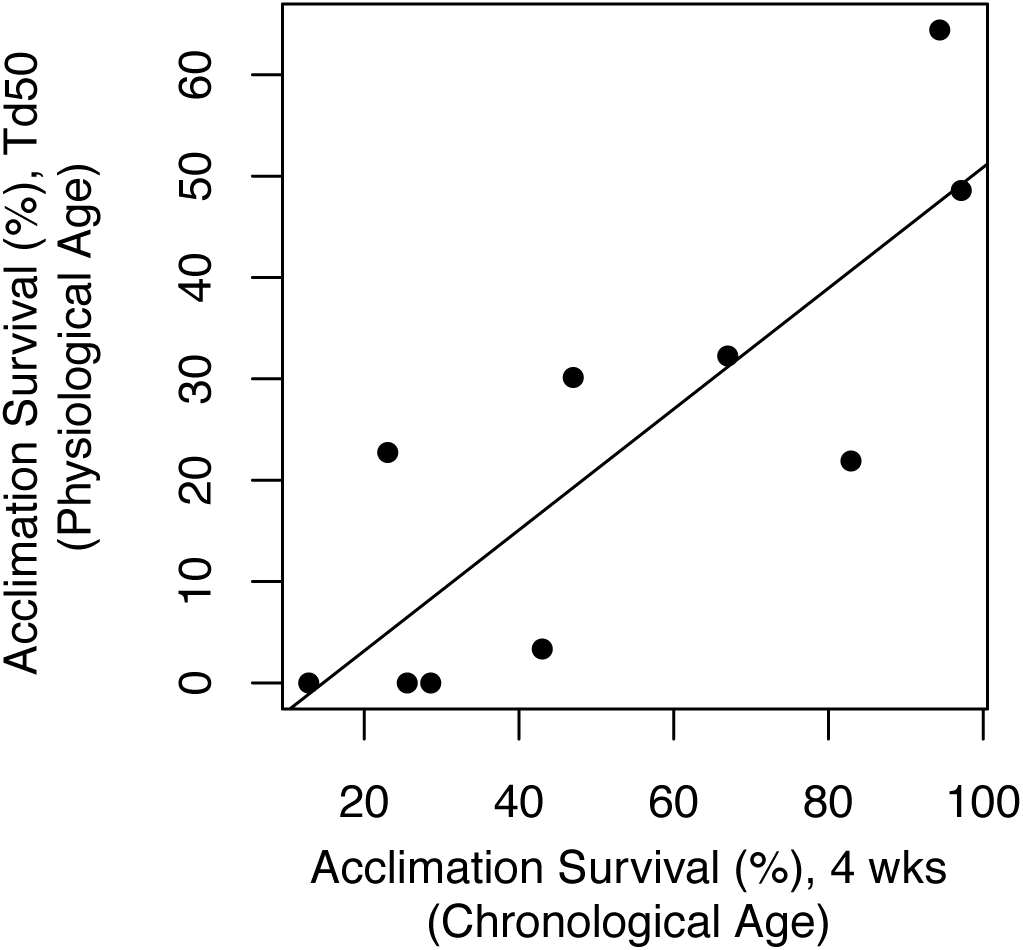
Stress tolerance of 4-week-old flies was correlated with stress tolerance of flies that were a similar physiological age (Td50). Acclimation and non-acclimation survivorship was measured for 10 DGRP lines that were aged to four weeks (chronological age) and until populations in experimental vials reached 50% of the starting population (physiological age). Physiologically-aged flies did not survive the non-acclimation treatment. Acclimation survivorship in four-week old flies is a good predictor of acclimation survivorship in flies that are approximately the same physiological age (R = 0.833, *P* < 0.003). Points shown indicate average acclimation survival for the 10 DGRP lines.

### Genetic architecture

In one-week-old flies, association mapping identified 24 polymorphisms and 23 genes associated with acclimation survival, 22 polymorphisms and 14 genes associated with non-acclimation survival, 45 polymorphisms and 23 genes associated with acclimation score, and 20 polymorphisms and 9 genes associated with starvation resistance (Table 1). In four-week-old flies, association mapping identified 31 polymorphisms and 28 genes associated with acclimation survival, 69 polymorphisms and 48 genes associated with non-acclimation survival, 26 polymorphisms and 6 genes associated with acclimation score, and 27 polymorphisms and 22 genes associated with starvation resistance (Table 1). Surprisingly, no polymorphisms or genes were shared within phenotype across age or between phenotypes (Fig. S2; Table S5).

Several polymorphisms were associated with genes that have been previously associated with cold-, starvation-, or age-related phenotypes, and were distributed across the phenotypes measured in this study (Table S5 and references therein). Out of all genes identified in our study (Table 1, Table S5), 54 have been previously associated with cold acclimation or with a cold-sensitive phenotype in *Drosophila,* 18 have been previously associated with starvation response or stress, and 59 have been previously associated with aging or lifespan. For example, *Cht2,* involved in chitin binding, has been previously associated with cold acclimation response (MacMillan et al., 2016) and was associated with four-week starvation resistance in this study. *Meltrin,* associated with one-week starvation resistance in our study, has been previously associated with cold acclimation response and age-specific fitness (Durham et al., 2014; MacMillan et al., 2016). *CG10916,* associated with four-week non-acclimation survival in our study, has previously been associated with determination of adult lifespan as well as cold acclimation (MacMillan et al., 2016; Paik et al., 2012; Vermeulen et al., 2013). Several genes (28) were also associated with oxidative stress resistance, which has been associated with aging and senescence (Schwarze et al., 1998). For example, *decay, rg,* and *Pde1c* have been previously associated with oxidative stress and were associated with four-week acclimation survival or one-week non-acclimation survival in our study (Table S5). Additional details describing the function of each gene and associated references are listed in Table S5. Despite the previous reporting of genes that are associated with aging or stress phenotypes, no gene ontology (GO) categories were overrepresented following enrichment analysis,.

**Table 1.** Quantitative genetic estimates (± S.E.) for all phenotypes as they vary with age and the number of polymorphisms (generalized as SNPs) and genes significantly associated with each phenotype identified by GWAS with a threshold of −log10(5). All heritabilities reported are broad-sense and are greater than 0.

| Phenotype | No.<br>Lines | Age<br>(weeks) | Mean | H<sup>2</sup> | CV<sub>G</sub> | CV<sub>E</sub> | No.<br>SNPs | No.<br>Genes |
| --- | --- | --- | --- | --- | --- | --- | --- | --- |
| Acclimation | 101 | 1 | 63.4 (1.1) | 0.26 (0.04) | 22.68 | 38.58 | 24 | 23 |
| Acclimation | 101 | 4 | 55.2 (1.2) | 0.19 (0.04) | 25.60 | 52.70 | 31 | 28 |
| Non-acclimation | 101 | 1 | 30.4 (1.3) | 0.33 (0.04) | 58.76 | 84.52 | 22 | 14 |
| Non-acclimation | 101 | 4 | 14.1(0.9) | 0.26 (0.05) | 83.86 | 141.09 | 69 | 48 |
| Acclimation Score | 101 | 1 | 33.03 (1.4) | 0.20 (0.03) | 50.53 | 101.67 | 45 | 23 |
| Acclimation Score | 101 | 4 | 41.1 (1.3) | 0.14 (0.02) | 31.94 | 79.03 | 26 | 6 |
| Starvation | 164 | 1 | 51.89 (0.4) | 0.34 (0.03) | 13.51 | 18.66 | 20 | 9 |
| Starvation | 164 | 4 | 44.62 (0.5) | 0.13 (0.02) | 14.00 | 36.45 | 27 | 22 |

### Evolutionary theories of aging and the decline in stress response

#### Shifting genetic architecture within phenotypes across age

Each phenotype was associated with a unique set of polymorphisms across age and among phenotypes (Table 1, Fig. S2, Table S5). Thus, the lack of overlap of associated polymorphisms across age within phenotypes suggests the genetic architecture shifted and that genetic control of the phenotypes was age-specific. MA not only predicts that associated polymorphisms at each age are unique, but also that associated polymorphisms have positive additive effects in young individuals that remain positive or approach 0 in older individuals. Conversely, AP predicts associated polymorphisms with positive additive effects in young individuals will have negative additive effects in old individuals. We calculated the additive effects of all significantly associated polymorphisms for the stress response phenotypes that declined with age (acclimation survivorship, non-acclimation survivorship, and starvation resistance; Fig. 4) for the one- and four-week response. When additive effects of the associated polymorphisms in one-week-old flies were calculated for the same phenotype in four-week-old flies, the additive effects were either closer to 0 or of the same sign (i.e. not antagonistic; Fig. 4; Table S6). The reverse comparison resulted in the same pattern; for example, additive effects of four-week acclimation survival polymorphisms on one-week acclimation survival were smaller and closer to 0 than the additive effects of four-week acclimation survival polymorphisms on four-week acclimation survival (Fig. 4; Table S6). This pattern of unique associated polymorphisms and additive effects that decrease with age was observed for each of the phenotypes that declined with age (starvation resistance, acclimation survival, and non-acclimation survival) and is consistent with MA.

**Figure 4.**
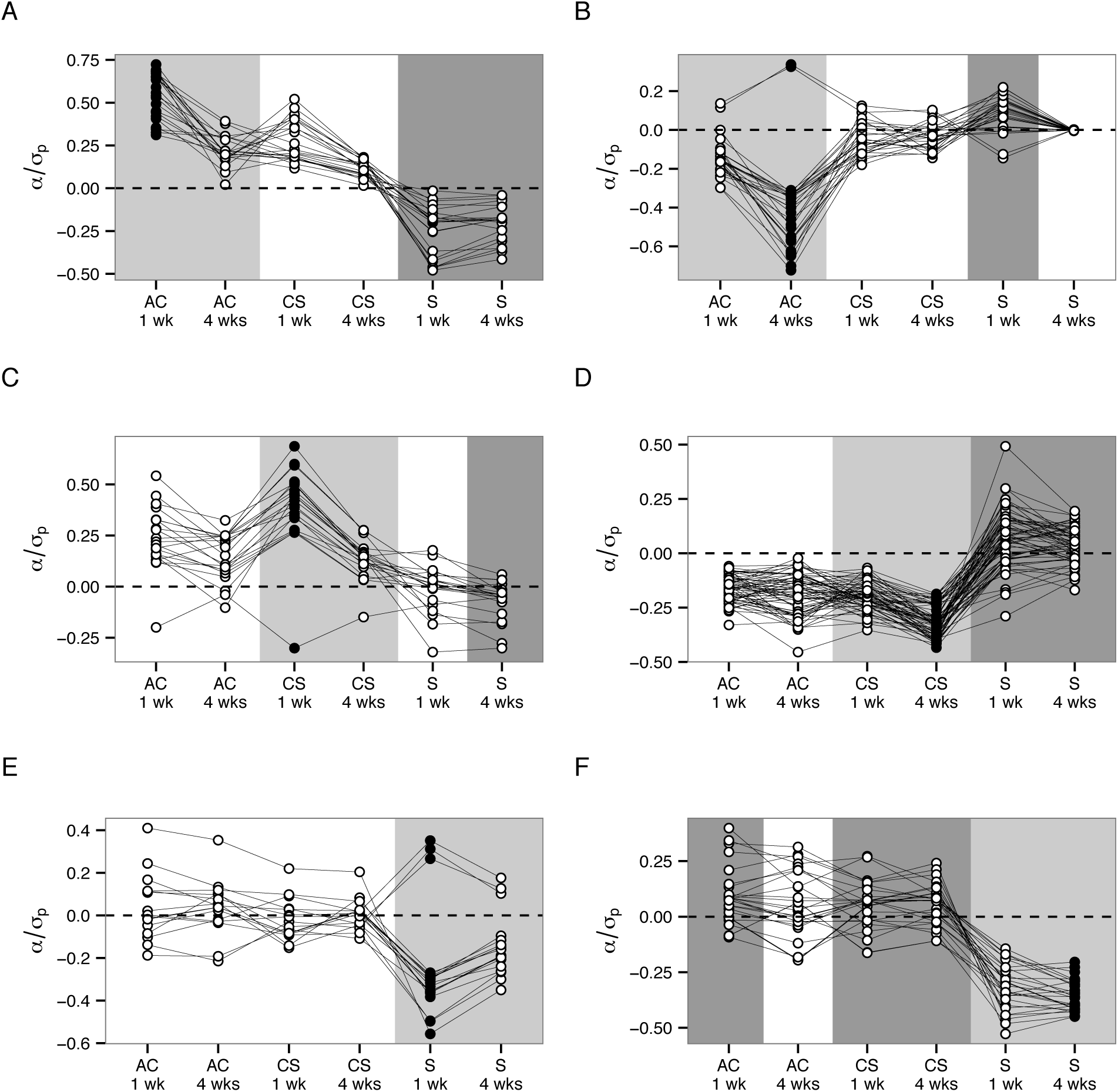
Standardized additive effects (α/σ_p_) of polymorphisms associated with each phenotype across age (A – F). In each plot, black solid points indicate which age and phenotype the polymorphisms were significantly associated with. Open points indicate the calculated additive effects of those polymorphisms on each of the other phenotypes. The light grey shading highlights the ‘within phenotype’ change in the additive effect of polymorphisms across age. Darker grey shading highlights ‘between phenotype’ comparisons where antagonistic effects of polymorphisms were found based on analysis of 95% confidence interval around the average additive effect (Table S6). Unshaded sections of each plot indicate calculated additive effects that are consistent with MA relative to the additive effects of the polymorphisms on their associated phenotype. In each plot, AC indicates acclimation survival, CS indicates non-acclimation survival, and S indicates starvation resistance. A. Polymorphisms associated with acclimation survival of one-week-old flies all had positive additive effects on the phenotype that became smaller when the additive effects were calculated for four-week-old acclimation survival and non-acclimation survival of one- and four-week-old flies. Polymorphisms had antagonistic effects on starvation resistance at both ages. B. Polymorphisms associated with acclimation survival of four-week-old flies had smaller additive effects on every other phenotype except one-week starvation resistance, where the effects were largely antagonistic. C. Polymorphisms associated with non-acclimation survival of one-week-old flies had smaller additive effects on every other phenotype except four-week starvation resistance, where the effects were largely antagonistic. D. Polymorphisms associated with non-acclimation survival of four-week-old flies had smaller additive effects on every other phenotype except starvation resistance at both ages, where the effects were largely antagonistic. E. Polymorphisms associated with one-week starvation resistance had smaller additive effects on every other phenotype. F. Polymorphisms associated with four-week starvation resistance had antagonistic effects on every other phenotype except acclimation survival in four-week-old flies and starvation resistance in one-week-old flies.

MA also predicts that, for phenotypes that decline with age, the coefficient of genetic variance (CV_G_) will increase with age and that the genetic correlation between the phenotype in young individuals will be non-negative (either not different from 0 or positively correlated) with the phenotype in old individuals. While AP does not have predictions specific to the coefficient of genetic variance (Houle et al., 1994; Partridge and Barton, 1993), AP does predict a negative genetic correlation between the phenotype in young individuals and in old individuals. Variance component analysis provided further support for MA for each phenotype. We observed increase in CV_G_ for each phenotype, although the increase was slight for starvation resistance (CV_G,young_ = 13.51 versus CV_G, old_ 14; Table 1). Also consistent with MA predictions, genetic correlations for each phenotype across age were either significantly positive (acclimation survival: R_G_ = 0.70 ± 0.3 S.E.; starvation resistance: R_G_ = 0.69 ± 0.3 S.E.), or not statistically different from 0 (non-acclimation survival: R_G_ = 0.43 ± 0.3 S.E.; Table S8). The lack of negative genetic correlations for each phenotype between young and old individuals combined with the increase in the coefficient of genetic variance with age provides additional evidence that supports the MA mechanism.

#### Shifting genetic architecture between phenotypes with age

Comparisons of associated polymorphisms for each phenotype measured in this study demonstrated unique genetic control for each phenotype across age. However, the lack of overlap in associated polymorphisms does not necessarily mean that the polymorphisms detected for each phenotype do not influence variation in other phenotypes at other ages, but rather that the additive effect was too small to be detected by association mapping. Polymorphisms associated with one phenotype (e.g. one-week acclimation survival) may have small but important additive effects on other phenotypes (e.g. four-week starvation resistance), and if this is the case, the interpretation is similar to the comparison of additive effects within phenotypes across age reported above (e.g. Fig. 1). If the additive effect is close to 0 and of the same sign, this supports the MA mechanism; if the additive effect is different from 0 and of the opposite sign, this supports the AP mechanism (Fig. 1).

To test for the presence of pleiotropic effects of associated polymorphisms on other phenotypes measured in this study, we calculated the average standardized additive effect of associated polymorphisms for each phenotype on every other phenotype and age (Fig. 4; Table S6). For example, the additive effects of the set of associated polymorphisms with acclimation survival in one-week-old flies were calculated for one- and four-week non-acclimation survival and one- and four-week starvation resistance (Fig. 4A). Confidence intervals were used to determine if the calculated average additive effects were different from 0 (Table S6).

Approximately half of the calculated average effects were not different from 0 (55.6%), and 27.8% of the comparisons resulted in average effects with a sign opposite that of the average effect of the polymorphisms in the phenotype with which they were significantly associated (suggesting an antagonistic relationship; Table S6). The antagonistic effects of polymorphisms between phenotypes were often, but not always, reciprocal (Fig. 4). For example, polymorphisms associated with one-week acclimation survival had an average antagonistic additive effect on starvation resistance at both ages (Fig. 4A), while polymorphisms associated with four-week starvation resistance had an average antagonistic additive effect on only one-week acclimation survival (Fig. 4F). In an aging context, evidence from additive effect comparisons support both AP and MA across age, depending on the phenotypes being compared; however, MA was more common based on apparent independence of additive effects (average additive effects were not different from 0).

Phenotypes that appeared to be antagonistically pleiotropic such that polymorphisms increased the phenotype in young flies but decreased the phenotype in old flies include acclimation survival and starvation resistance (Fig. 4A, B, F) and non-acclimation survival and starvation resistance (Fig. 4C, D, F). Polymorphisms associated with one-week starvation resistance did not have an antagonistic effect on average on any other phenotype, although several individual polymorphisms did have antagonistic effects on other phenotypes (Fig. 4E; Table S6). The antagonistic relationship between one-week acclimation survival and four-week starvation resistance was further supported by a significant negative genetic correlation between traits across ages (R_G_ = −0.47 ± 0.2 S.E.; Table S8). However, all other combinations of phenotypes involved effects and genetic correlations that were not different from 0 (Fig. 4; Table S6, S8), and were thus more consistent with the predictions of MA.

#### Evidence of selection and phenotypic trade-offs

With the exception of acclimation score, the additive effects of the majority of polymorphisms significantly associated with the phenotypes measured in our study were of the same sign (i.e. most additive effects were positive or negative; Fig. 4 and 5, Table S5). To determine if more additive effects of positive sign were associated with the phenotype than expected by chance, we used the QTLST to test for evidence of selection. More positive additive effects than expected by chance were observed for one-week acclimation survival, non-acclimation survival, and acclimation score suggesting selection increased these phenotypes in the founding population of the DGRP (Fig. 4 and Fig. 5A, C, E). The QTLST was also significant for four-week acclimation survival, non-acclimation survival, and both one- and four-week starvation resistance, suggesting selection has acted to decrease these phenotypes (Fig. 4 and Fig. 5B, D, G, H). However, because the effectiveness of natural selection is expected to decline with age, this significant result is likely the result of a correlated response resulting from selection on young phenotypes. The signs of the additive effects of polymorphisms associated with acclimation score in four-week-old flies were more mixed than any other phenotype, leading to a non-significant QTLST for this phenotype (Fig. 4 and Fig. 5F).

**Figure 5.**
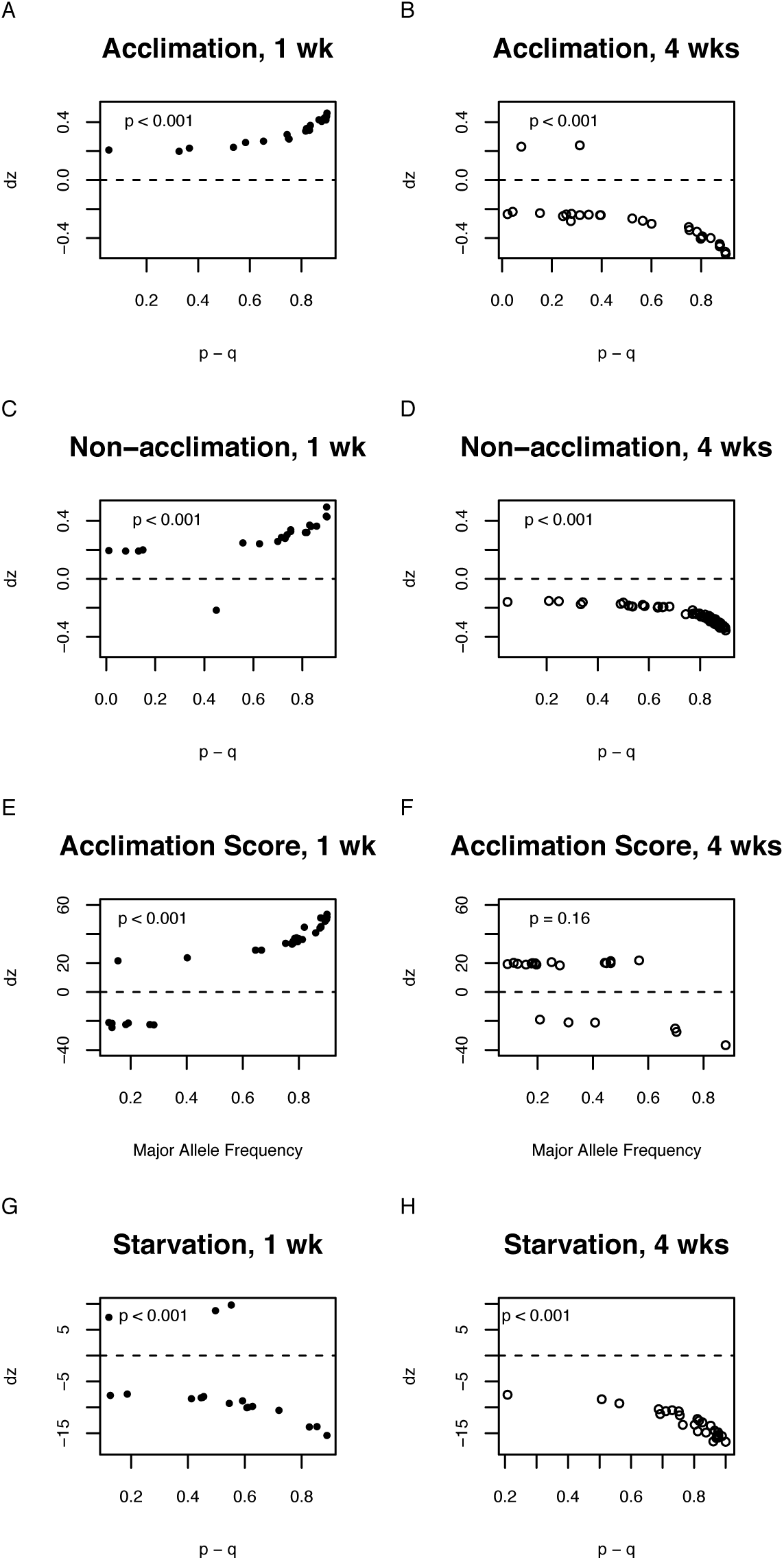
Plots of allele frequency (p – q) against the effect of significant polymorphisms reflected as change in phenotype (dz). Inset numbers are QTLST probabilities that directional selection influenced the phenotype.

## Discussion

### Genetic variation in age-specific decline in stress tolerance

As expected, the average phenotypic responses for most phenotypes declined with age, with the exception of plasticity measured as acclimation score (Fig. 2). For each phenotype, we observed significant genetic variation across ages, with some genotypes exhibiting increased stress resistance with age. We investigated variation in longevity among DGRP lines by comparing cold tolerance responses measured at four weeks to those in physiologically aged flies at the point when the population reached Td50. We know that lifespan for virgin female flies varies from approximately 20 days to approximately 80 days in the DGRP (Ivanov et al., 2015). This variation could result in an inequivilence of line comparisons at four weeks. Based on (Ivanov et al., 2015) longevity estimates, some lines may be only half way through their life span while others may be closer to the end of their life span at four weeks of age. Furthermore, genetic shifts known to be associated with senescence would be further advanced in lines that age more quickly (Pletcher et al., 2002). The significant positive correlation in acclimation survival between chronologically aged (four-week) flies and physiologically aged (Td50) flies suggests that, despite variation in the rate of senescence among lines, the four-week point is representative of how an “old” fly responds to cold stress. Additionally, this subset of the DGRP lines reached Td50 well after four weeks of age (Table S4), indicating that the four-week aging period does not lead to many lines entering a late-life plateau in mortality and that our lines were likely in the aging phase (Charlesworth, 2001; Curtsinger, 2016)

The only phenotype in our study that behaved unexpectedly and did not decline with age was plasticity measured as acclimation score. Acclimation score was significantly higher in four-week-old flies (Fig. 2G), where we expected this phenotype to remain constant across age. The age-related response in acclimation score was driven by the male response. Four-week-old male flies had a stronger age-related decline in non-acclimation survival compared to acclimation survival. Therefore, the observed increase in acclimation score for our population has at least two interpretations. First, plasticity at the population level may increase with age, potentially as a compensatory mechanism to overcome the overall loss of basal cold tolerance (Fig. 2F). Throughout the season, natural populations of *D. melanogaster* are expected to be composed of increasingly old individuals such that by the time temperatures begin to cool at the beginning of the fall season, a greater proportion of populations is composed of older individuals (Behrman et al., 2015). If older individuals are less cold tolerant, they may still be able to tolerate cold temperature exposures through increased capacity for adaptive plasticity through acclimation. Second, acclimation pretreatment appears to have had a less detrimental effect on survival in four-week-old flies compared to one-week-old flies (Fig. 2I). In one-week-old flies, acclimation improved survival in the majority of lines; however, several lines (14%) had negative acclimation scores, indicating that exposure to 4°C prior to the −6°C exposure was more damaging than the −6°C exposure alone (Fig. 2I; Gerken et al., 2015). Only 2% of lines tested had negative acclimation scores at four weeks suggesting that acclimation may be more likely to either improve or not affect survival following cold stress at four weeks. Shifts in the effect of acclimation on survival were observed when genotypes were reared under different developmental conditions or experienced altered thermal regimes (Gerken et al., 2015; Kelty and Lee, 2001); our results suggest that the age of the individual has an important influence on the effect of acclimation on survival as well.

### MA describes age-related change within individual phenotypes

When each phenotype was considered separately across age, we found support for MA, satisfying predictions based on analysis of quantitative genetic parameters (Charlesworth, 2001; Charlesworth and Hughes, 1996; Hughes et al., 2002; Leips et al., 2006; Reynolds et al., 2007). First, the coefficient of genetic variance increased with age for each phenotype (Table 1). The increase in CVG in starvation resistance was less drastic than other phenotypes, but when sexes were analyzed separately, the increase was more dramatic (Table S7). An increase in CV_G_ indicates that a greater proportion of the phenotypic variance can be explained genetically at four weeks of age (Charlesworth, 2001; Charlesworth and Hughes, 1996; Houle et al., 1994) and this increase is consistent with the hypothesis that the age-related decline in the phenotype is the result of the accumulation of deleterious age-specific polymorphisms that influence variation in the phenotype (Charlesworth, 2001; Charlesworth and Hughes, 1996; Engström et al., 1989).

Second, MA predicts that a phenotype is controlled by unique sets of genes across age (Charlesworth, 2001, 1994; Charlesworth and Hughes, 1996; Maklakov et al., 2015; Medawar, 1952; Partridge and Barton, 1993; Rose, 1991). We detected unique sets of polymorphisms that were associated with each phenotype across age. This is consistent with age-specific association patterns presented by (Durham et al., 2014) who determined age-related change in fecundity was also influenced by MA. The genetic independence of phenotypes across age was further supported by the non-significant or significantly positive genetic correlations for each phenotype across age (Table S8). While positive genetic correlations suggest that the genetic control of the phenotype across age is not independent (inconsistent with MA), Charlesworth (2001) and Maklakov et al. (2015) suggest that positive pleiotropy can lead to significantly positive genetic correlations across age under MA.

Positive pleiotropy (Maklakov et al., 2015) expands on MA predictions in that, instead of limiting polymorphisms to very narrow windows of effect, polymorphisms can influence a wider window of ages but with lower additive effects (Maklakov et al., 2015). Thus, the positive genetic correlations across age are the result of the associated polymorphisms having a slightly wider window of age-specific effects that ultimately influence the phenotype at other ages. This pattern was observed for acclimation and non-acclimation survival and starvation resistance across age; for all phenotypes, the four-week associated polymorphisms had negative additive effects on the one-week phenotype that were smaller than the effect of the polymorphisms on the four-week phenotype. This suggests that four-week polymorphisms that led to decline in each phenotype do have small pleiotropic effects (in the same direction) at one week of age. The failure of natural selection to remove polymorphisms that have small negative effects early in life and larger negative effects later in life is one way in which senescence can evolve and is consistent with expectations of positive pleiotropy under MA (Maklakov et al., 2015; Wachter et al., 2014, 2013).

Our third piece of evidence to support MA comes from our novel approach of calculating the additive effects of polymorphisms across age. We calculated the additive effects of polymorphisms associated with the one-week phenotype in the four-week phenotype data (and vice versa). Under MA, we expected the additive effect of the one-week polymorphisms to be smaller and in the same direction (i.e. they have the same sign) when the additive effects were calculated for the four-week response data (Maklakov et al., 2015). If the sign of a particular polymorphism had flipped (a polymorphism with a positive effect in one age had a negative effect in the other age), this would have suggested that an antagonistic relationship existed and would have supported AP (Maklakov et al., 2015). For all phenotypes, the calculated additive effects of age-specific polymorphisms across age were either closer to 0 and/or in the same direction, providing definitive support for the role of MA in the age-related decline in the stress responses measured (Fig. 4).

### MA and AP describe age-related variation between phenotypes

Our novel extension of association mapping through the calculation of additive effects of polymorphisms across phenotypes allowed us to investigate the role MA and AP on age-specific responses between phenotypes as well. Though each phenotype and age was associated with a unique set of polymorphisms (consistent with predictions for MA), we found support for AP between several phenotypes (Fig. 4). We observed a significantly negative phenotypic correlation between one-week acclimation survival and both one- and four-week starvation resistance, corroborating a pattern reported by (Hoffmann et al., 2005) (Table S8). We also observed a significantly negative genetic correlation between one-week acclimation survival and four-week starvation resistance (Table S8), suggesting that AP (between one-week acclimation survival and four-week starvation resistance) influenced age-related change in these phenotypes. When the additive effects of polymorphisms associated with four-week starvation were calculated for both one-week cold tolerance phenotypes (Table S6), all four-week starvation resistance associated polymorphisms, which had negative additive effects on four-week starvation resistance, had positive additive effects on one-week acclimation survival and one-week non-acclimation survival (Fig. 4F). The change in sign of the additive effects of four-week starvation resistance associated polymorphisms on both of the one-week cold tolerance phenotypes is strong evidence to support the role of AP in age-related decline in starvation resistance.

Additional examples of AP existed between phenotypes in our data as well and were identified through confidence interval analysis of additive effects (Fig. 4; Table S6). Specifically, one-week acclimation and non-acclimation survival polymorphisms had positive effects on their respective phenotypes across age but negative additive effects on four-week starvation resistance (Fig. 4A – D; Table S6). This pattern is consistent with that discussed above and again suggests that age-related change in starvation resistance is influenced by AP with cold tolerance. Interestingly, four-week acclimation and non-acclimation survival polymorphisms had largely positive additive effects on one-week starvation resistance (Fig. 4B, D; Table S6). This pattern suggests that AP may also be contributing to age-related decline in acclimation and non-acclimation survival. With evidence from our examination of acclimation and non-acclimation survival across age (discussed above), and the apparent role of MA and positive pleiotropy for age-related change within these phenotypes, it is evident that it may not be possible to fully disentangle the roles of MA and AP on the age-related decline in phenotypes. In essence, polymorphisms that increase one phenotype in young individuals and decrease another phenotype in old individuals through AP may also contribute to age-related change within the phenotype through positive pleiotropy under MA. These results demonstrate the need for caution in interpreting the lack of overlap in significant associated polymorphisms as support for MA in isolation of other evidence because AP may still be playing an important role in the age-related change in phenotypes.

We also found support for the role of MA between phenotypes across age. Age-related change in acclimation survival and non-acclimation survival appears to be evolving largely independently of the other phenotype under MA as we found either non-significant or significantly positive phenotypic and genetic correlations between all age combinations of these phenotypes (Table S6, S8). The calculated additive effects of polymorphisms between phenotypes and ages uphold this interpretation to a large degree as well (Fig. 4A – D; Table S6). One-week acclimation survival polymorphisms, which had positive effects on one-week acclimation survival, all had positive additive effects when calculated for both one- and four-week non-acclimation survival (Fig. 4A). On average, additive effects of polymorphisms associated with four-week acclimation survival had additive effects that were not different from zero, although some individual polymorphisms did have antagonistic effects on one- and four-week non-acclimation survival. All but two polymorphisms associated with one-week non-acclimation survival had additive effects of the same sign on one- and four-week acclimation survival, and all polymorphisms associated with four-week non-acclimation survival had additive effects of the same sign on one- and four-week acclimation survival. While the small number of individual polymorphisms with antagonistic additive effects across phenotype may impact age-related change in these cold tolerance phenotypes, it likely that this impact is small in comparison to the role of MA and positive pleiotropy.

### Natural selection shapes phenotypic variation across age

Our data recapitulate previously reported relationships between different measures of cold tolerance and starvation resistance (Table S8; Gerken et al., 2015; Hoffmann et al., 2005; Sinclair and Roberts, 2005); however, we have added another layer to these relationships by considering the influence of age and natural selection on these phenotypes. Specifically, the direction of the additive effects of polymorphisms associated with each phenotype may reflect the role natural selection has played in the evolution of the phenotype. Through applying a basic sign test (QTLST probabilities) to the additive effects of the associated age-specific polymorphisms, it is evident that the signs of the effects are not randomly associated with the frequency of the allele (Fig 5A – E, G, H). If a phenotype is evolving neutrally, polymorphisms that influence the phenotype are equally likely to have positive or negative effects (Anderson and Slatkin, 2003; Orr, 1998; Rice and Townsend, 2012). Signatures of directional selection are detected when this null hypothesis is rejected (Anderson and Slatkin, 2003; Muir et al., 2014; Orr, 1998; Rice and Townsend, 2012; Rieseberg et al., 2002). Because we observed an overabundance of major alleles with positive effects on one-week acclimation and non-acclimation survival, this suggests that natural selection favored polymorphisms that increase cold tolerance phenotypes in the population from which the DGRP was established (Fig. 4A, C and 5A, C; Table S5). This finding is consistent with evidence of selection for cold tolerance in natural populations of *D. melanogaster* (Bergland et al., 2014), as well as previous reports of majority positive additive effects of polymorphisms associated with chill coma recovery (Mackay et al., 2012).

Conversely, most of the polymorphisms associated with cold tolerance at four weeks of age were negative (Fig. 4B, D), indicating that the major alleles decreased survival following cold stress. While the QTLST was significant for these late-acting polymorphisms, it is very unlikely that natural selection directly led to this pattern. Instead, polymorphisms associated with acclimation and non-acclimation survival in old individuals likely arose through mutation and were maintained in the population because their negative effect on survival in young individuals was small relative to the four-week additive effects (Charlesworth, 2001; Houle et al., 1994; Maklakov et al., 2015)Fig. 4B, D). Similarly, in both young and old individuals, most of the additive effects of starvation resistance associated polymorphisms were negative (Fig. 4E, F). This pattern suggests that age-related change in starvation resistance is influenced by positive selection on acclimation and non-acclimation survival in young individuals. Alternatively, the effectiveness of natural selection on starvation resistance may be constrained by pleiotropy between one-week acclimation survival and one-week starvation resistance (many one-week acclimation and non-acclimation polymorphisms had negative effects on one-week starvation resistance; Fig. 4A, C). Some positive selection on starvation resistance in young individuals may provide a mechanism for the retention of four-week acclimation and non-acclimation survival associated polymorphisms that have increasingly negative effects with age.

### Implications for evolutionary theories of aging

Efforts to more fully understand and test evolutionary theories of aging have encouraged expansion and clarification of predictions of both MA and AP mechanisms (Charlesworth, 2001; Houle et al., 1994; Maklakov et al., 2015; Reynolds et al., 2007; Wachter et al., 2014, 2013). Our novel extension of existing methods not only revealed the relative importance of MA and AP for age-related change in stress response, but also verified recent hypotheses that present expansions on the theory of MA. When originally formulated, the MA mechanism predicted that fitness was controlled by polymorphisms that had very narrow windows of effect (Charlesworth and Hughes, 1996; Medawar, 1952; Rose, 1991), but evidence from several studies has indicated that it is more likely that polymorphisms which contribute to late-life decline in fitness and age-related change in phenotypes have wide windows and increasingly large effects across age (Houle et al., 1994; Maklakov et al., 2015). Patterns of additive effects of associated polymorphisms identified in our study align well these elaborations of MA predictions and provide empirical support for positive pleiotropy. In addition, we provide evidence supporting the potential for positive genetic correlations between young and old phenotypes under MA, demonstrating through patterns of additive effects of polymorphisms that, while the size of the effect may change across age, the direction of the effect is often preserved (Maklakov et al., 2015; Reynolds et al., 2007).

Our use of calculated additive effects also allowed us to overcome difficulties in characterizing the role of AP in age-related change in phenotypes. The isolated analysis of the coefficient of genetic variance and genetic correlation is problematic because the signature of AP may be too small to detect due to near fixation of segregating alleles at the antagonistic loci (Moorad and Promislow, 2009; Partridge and Barton, 1993; Schnebel and Grossfield, 1988). For this reason, association mapping may be particularly biased against the detection of AP loci. However, by comparing the effects of polymorphisms calculated for each age and phenotype pair, we are able to overcome this bias against polymorphisms that have small effects.

Very few studies present convincing evidence of the influence of both MA and AP on age-related change within and among phenotypes (but see Leips et al., 2006), but we have demonstrated that both mechanisms contribute to age-related change in stress response. It is clear from our results that individual polymorphisms that are significantly associated with phenotypes at different ages can contribute to age-related decline within and among phenotypes in patterns that are consistent with both MA and AP. Thus, the evolution of senescence and associated decline in fitness is influenced by a combination of natural selection acting on correlated phenotypes that have non-independent antagonistic genetic architectures, as well as the accumulation of polymorphisms with negative effects that strengthen with age. It is likely that similar patterns will be observed for other phenotypes related to fitness as well, adding to our understanding of how evolution of aging and multivariate evolution are tightly intertwined.

## Acknowledgments

We thank Jennifer L. Delzeit, Olivia Eller, Mariah Brown, and Paul Klawinski for assistance with experimental fly set up and maintenance. We also thank Michael Tobler, Phil Freda, and Kate Jordan for comments on earlier versions of this manuscript. The National Science Foundation awards NSF1051770 and NSF1156571 to T.J.M supported this work.

## Author Contributions

E.R.E and T.J.M. conceived the experimental framework and prepared the manuscript. E.R.E. performed the research and the statistical analyses.

---

**Figure S1.**
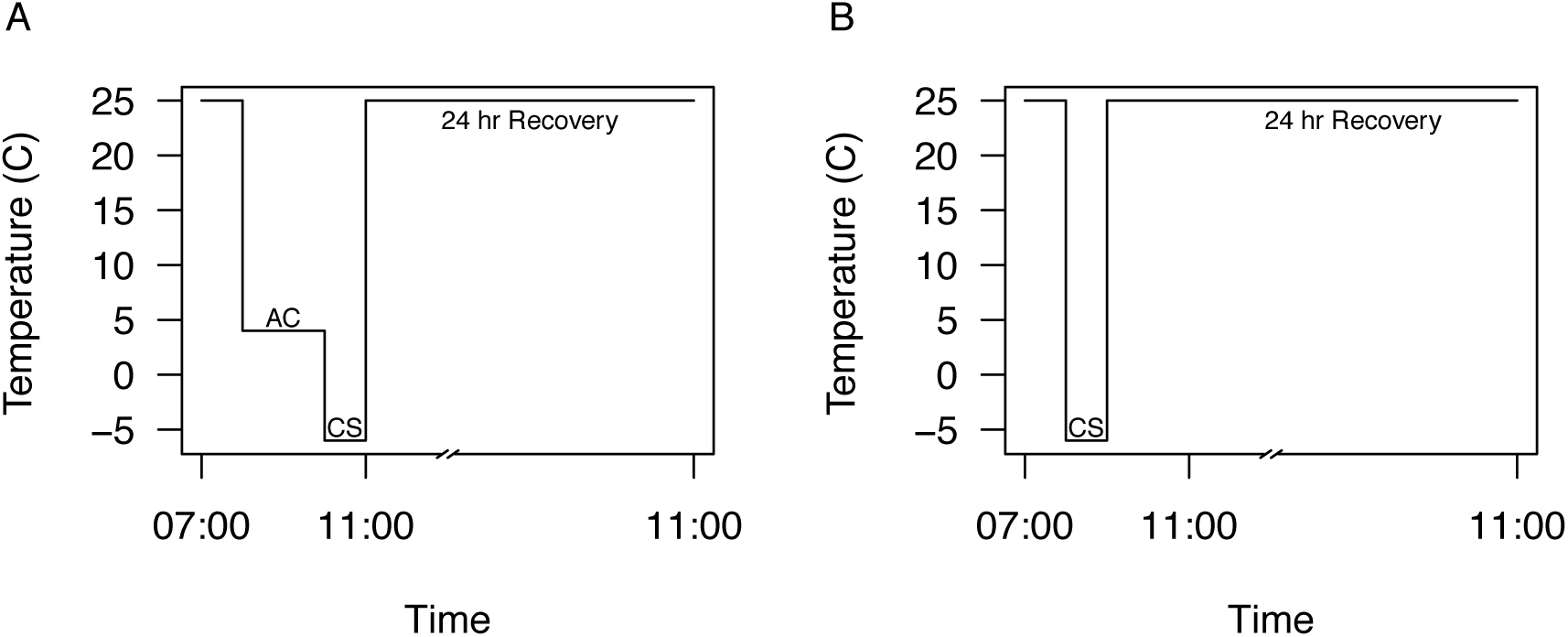
Graphical representation of acclimation (A) and non-acclimation treatment (B). Flies were maintained at 25°C during rearing and recovery, and lights on occurred at 07:00hrs. A. Flies were transferred from 25°C to 4°C for two hours for the acclimation (AC) treatment and then were transferred immediately to −6°C for one hour for the cold shock treatment (CS). Flies were placed on fresh media and allowed to recover for 24 hours at 25°C. B. Flies were transferred to −6°C for one hour for the cold shock treatment (CS) and were allowed to recover at 25°C for 24 hours on fresh media

**Figure S2.**
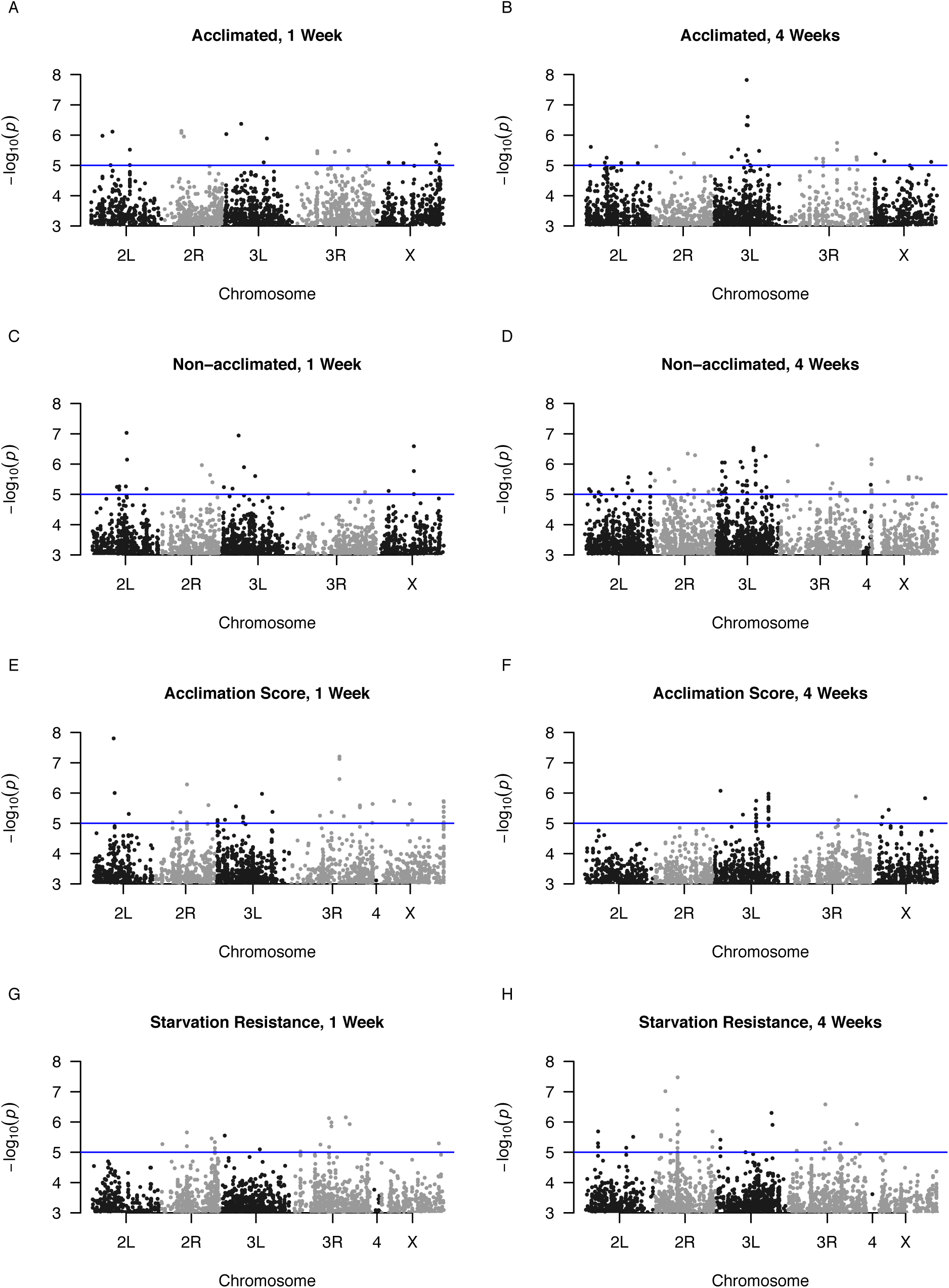
Manhattan plots of each phenotype. The significance threshold is indicated by the horizontal line at −log(5).

## References

Anderson, E.C., Slatkin, M., 2003. Orr’s quantitative trait loci sign test under conditions of trait ascertainment. Genetics 165, 445–446.

Behrman, E.L., Watson, S.S., O’Brien, K.R., Heschel, M.S., Schmidt, P.S., 2015. Seasonal variation in life history traits in two *Drosophila* species. J. Evol. Biol. 28, 1691–1704. doi: 10.1111/jeb.12690

Bergland, A.O., Behrman, E.L., O’Brien, K.R., Schmidt, P.S., Petrov, D.A., 2014. Genomic evidence of rapid and stable adaptive oscillations over seasonal time scales in *Drosophila*. PLoS Genet. 10, e1004775. doi:10.1371/journal.pgen.1004775

Bowler, K., Terblanche, J.S., 2008. Insect thermal tolerance: what is the role of ontogeny, ageing and senescence? Biol. Rev. 83, 339–355. doi:10.1111/j.1469-185X.2008.00046.x

Bozinovic, F., Calosi, P., Spicer, J.I., 2011. Physiological Correlates of Geographic Range in Animals. Annu. Rev. Ecol. Evol. Syst. 42, 155–179. doi:10.1146/annurev-ecolsys-102710-145055

Charlesworth, B., 2001. Patterns of age-specific means and genetic variances of mortality rates predicted by the mutation-accumulation theory of ageing. J. Theor. Biol. 210, 47–65.

Charlesworth, B., 1994. Evolution in Age-Structured Populations, 2nd ed. Cambridge University Press, New York, New York.

Charlesworth, B., Hughes, K.A., 1996. Age-specific inbreeding depression and components of genetic variance in relation to the evolution of senescence. Proc. Natl. Acad. Sci. 93, 6140–6145.

Colinet, H., Chertemps, T., Boulogne, I., Siaussat, D., 2015. Age-related decline of abiotic stress tolerance in young *Drosophila melanogaster* adults. Gerontol. Soc. Am. 00, 1–7.

Coulson, S.C., Bale, J.S., 1990. Characterisation and limitations of the rapid cold-hardening response in the housefly *Musca domestica* (Diptera: Muscidae). J. Insect Physiol. 36, 207–211.

Curtsinger, J.W., 2016. Retired flies, hidden plateaus, and the evolution of senescence in *Drosophila melanogaster*. Evolution 70, 1297–1306. doi:10.1111/evo.12946

Czajka, M.C., Lee, R.E., 1990. A rapid cold-hardening response protecting against cold shock injury in *Drosophila melanogaster*. J. Exp. Biol. 148, 245–254.

Durham, M.F., Magwire, M.M., Stone, E.A., Leips, J., 2014. Genome-wide analysis in *Drosophila* reveals age-specific effects of SNPs on fitness traits. Nat. Commun. 5. doi:10.1038/ncomms5338

Engström, G., Liljedahl, L.E., Rasmuson, M., Bjödrklund, T., 1989. Expression of genetic and environmental variation during ageing. Theor. Appl. Genet. 77, 119–122.

Falconer, D.S., Mackay, T.F.C., 1996. Quantitative Genetics, 4th ed. Longman Group Ltd., Essex, England.

Fallis, L.C., Fanara, J.J., Morgan, T.J., 2014. Developmental thermal plasticity among *Drosophila melanogaster* populations. J. Evol. Biol. 27, 557–564. doi:10.1111/jeb.12321

Fisher, R.A., 1930. The Genetical Theory of Natural Selection. Oxford University Press, Chicago.

Gerken, A.R., Eller, O.C., Hahn, D.A., Morgan, T.J., 2015. Constraints, independence, and evolution of thermal plasticity: Probing genetic architecture of long- and short-term thermal acclimation. Proc. Natl. Acad. Sci. 112, 4399–4404.

Haldane, J.B.S., 1941. New Paths in Genetics. Allen and Unwin, London.

Hamilton, W.D., 1966. The moulding of senescence by natural selection. J. Theor. Biol. 12, 12–45.

Hoffmann, A.A., Hallas, R., Anderson, A.R., Telonis-Scott, M., 2005. Evidence for a robust sex-specific trade-off between cold resistance and starvation resistance in Drosophila melanogaster. J. Evol. Biol. 18, 804–810. doi:10.1111/j.1420-9101.2004.00871.x

Hope, R. M., 2013. Rmisc: Ryan Miscellaneous.

Houle, D., Hughes, K.A., Hoffmaster, D.K., Ihara, J., Assimacopoulos, S., 1994. Effects of spontaneous mutation on quantitative traits. I. Variances and covariances of life history traits. Genetics 138, 773–785.

Huang, W., Massouras, A., Inoue, Y., Peiffer, J., Ramia, M., Tarone, A.M., Turlapati, L., Zichner, T., Zhu, D., Lyman, R.F., Magwire, M.M., Blankenburg, K., Carbone, M.A., Chang, K., Ellis, L.L., Fernandez, S., Han, Y., Highnam, G., Hjelmen, C.E., Jack, J.R., Javaid, M., Jayaseelan, J., Kalra, D., Lee, S., Lewis, L., Munidasa, M., Ongeri, F., Patel, S., Perales, L., Perez, A., Pu, L., Rollmann, S.M., Ruth, R., Saada, N., Warner, C., Williams, A., Wu, Y.-Q., Yamamoto, A., Zhang, Y., Zhu, Y., Anholt, R.R.H., Korbel, J.O., Mittelman, D., Muzny, D.M., Gibbs, R.A., Barbadilla, A., Johnston, J.S., Stone, E.A., Richards, S., Deplancke, B., Mackay, T.F.C., 2014. Natural variation in genome architecture among 20. *Drosophila melanogaster* Genetic Reference Panel lines. Genome Res. 24, 1193–1208. doi:10.1101/gr.171546.113

Huey, R.B., Kearney, M.R., Krockenberger, A., Holtum, J.A.M., Jess, M., Williams, S.E., 2012. Predicting organismal vulnerability to climate warming: roles of behaviour, physiology and adaptation. Philos. Trans. R. Soc. B Biol. Sci. 367, 1665–1679. doi:10.1098/rstb.2012.0005

Hughes, K.A., Alipaz, J.A., Drnevich, J.M., Reynolds, R.M., 2002. A test of evolutionary theories of aging. Proc. Natl. Acad. Sci. 99, 14286–14291.

Imasheva, A.G., Loeschcke, V., Zhivotovsky, L.A., Lazebny, O.E., 1998. Stress temperatures and quantitative variation in *Drosophila melanogaster*. Heredity 81, 246–253.

Ivanov, D.K., Escott-Price, V., Ziehm, M., Magwire, M.M., Mackay, T.F.C., Partridge, L., Thornton, J.M., 2015. Longevity GWAS Using the *Drosophila* Genetic Reference Panel. J. Gerontol. A. Biol. Sci. Med. Sci. 70, 1470–1478. doi:10.1093/gerona/glv047

Ju, R.-T., Xiao, Y.-Y., Li, B., 2011. Rapid cold hardening increases cold and chilling tolerances more than acclimation in the adults of the sycamore lace bug, *Corythucha ciliata* (Say) (Hemiptera: Tingidae). J. Insect Physiol. 57, 1577–1582. doi:10.1016/j.jinsphys.2011.08.012

Kelty, J., Lee, R.E., 2001. Rapid cold-hardening of *Drosophila melanogaster* (Diptera: Drosophilidae) during ecologically based thermoperiodic cycles. J. Exp. Biol. 204, 1659–1666.

Lee, R.E., Chen, C., Denlinger, D.L., 1987. A rapid cold-hardening process in insects. Science 238, 1415–1417.

Leips, J., Gilligan, P., Mackay, T.F.C., 2006. Quantitative trait loci with age-specific effects on fecundity in *Drosophila melanogaster*. Genetics 172, 1595–1605. doi:10.1534/genetics.105.048520

Lyne, R., Smith, R., Rutherford, K., Wakeling, M., Varley, A., Guillier, F., Janssens, H., Ji, W., Mclaren, P., North, P., Rana, D., Riley, T., Sullivan, J., Watkins, X., Woodbridge, M., Lilley, K., Russell, S., Ashburner, M., Mizuguchi, K., Micklem, G., 2007. FlyMine: an integrated database for *Drosophila* and *Anopheles* genomics. Genome Biol. 8, R129.

Mackay, T.F.C., Richards, S., Stone, E.A., Barbadilla, A., Ayroles, J.F., Zhu, D., Casillas, S., Han, Y., Magwire, M.M., Cridland, J.M., Richardson, M.F., Anholt, R.R.H., Barrón, M., Bess, C., Blankenburg, K.P., Carbone, M.A., Castellano, D., Chaboub, L., Duncan, L., Harris, Z., Javaid, M., Jayaseelan, J.C., Jhangiani, S.N., Jordan, K.W., Lara, F., Lawrence, F., Lee, S.L., Librado, P., Linheiro, R.S., Lyman, R.F., Mackey, A.J., Munidasa, M., Muzny, D.M., Nazareth, L., Newsham, I., Perales, L., Pu, L.-L., Qu, C., Ràmia, M., Reid, J.G., Rollmann, S.M., Rozas, J., Saada, N., Turlapati, L., Worley, K.C., Wu, Y.-Q., Yamamoto, A., Zhu, Y., Bergman, C.M., Thornton, K.R., Mittelman, D., Gibbs, R.A., 2012. The *Drosophila melanogaster* Genetic Reference Panel. Nature 482, 173–178. doi:10.1038/nature 10811

MacMillan, H.A., Knee, J.M., Dennis, A.B., Udaka, H., Marshall, K.E., Merritt, T.J.S., Sinclair, B.J., 2016. Cold acclimation wholly reorganizes the *Drosophila melanogaster* transcriptome and metabolome. Sci. Rep. 6, 28999. doi:10.1038/srep28999

Maklakov, A.A., Rowe, L., Friberg, U., 2015. Why organisms age: Evolution of senescence under positive pleiotropy? BioEssays 37, 802–807. doi:10.1002/bies.201500025

Medawar, P.B., 1952. An Unsolved Problem of Biology. London: H.K. Lewis

Moorad, J.A., Promislow, D.E.L., 2009. What can genetic variation tell us about the evolution of senescence? Proc. R. Soc. B Biol. Sci. 276, 2271–2278. doi:10.1098/rspb.2009.0183

Morgan, T.J., Mackay, T.F.C., 2006. Quantitative trait loci for thermotolerance phenotypes in *Drosophila melanogaster*. Heredity 96, 232–242.

Muir, C.D., Pease, J.B., Moyle, L.C., 2014. Quantitative genetic analysis indicates natural selection on leaf phenotypes across wild tomato species (*Solanum* sect. *Lycopersicon;* Solanaceae). Genetics 198, 1629–1643. doi:10.1534/genetics.114.169276

Niehaus, A.C., Angilletta, M.J., Sears, M.W., Franklin, C.E., Wilson, R.S., 2012. Predicting the physiological performance of ectotherms in fluctuating thermal environments. J. Exp. Biol 215, 694–701. doi:10.1242/jeb.058032

Orr, H.A., 1998. Testing natural selection vs. genetic drift in phenotypic evolution using quantitative trait locus data. Genetics 149, 2099–2104.

Overgaard, J., Sørensen, J.G., Jensen, L.T., Loeschcke, V., Kristensen, T.N., 2010. Field tests reveal genetic variation for performance at low temperatures in *Drosophila melanogaster*. Funct. Ecol. 24, 186–195.

Paik, D., Jang, Y.G., Lee, Y.E., Lee, Y.N., Yamamoto, R., Gee, H.Y., Yoo, S., Bae, E., Min, K.-J., Tatar, M., Park, J.-J., 2012. Misexpression screen delineates novel genes controlling *Drosophila* lifespan. Mech. Ageing Dev. 133, 234–245. doi:10.1016/j.mad.2012.02.001

Partridge, L., Barton, N.H., 1993. Optimality, mutation and the evolution of ageing. Nature 362, 305–311.

Phillips, P.C., 1998. H2boot: bootstrap estimates and tests of quantitative genetic data. University of Texas at Arlington.

Pletcher, S.D., Houle, D., Curtsinger, J.W., 1998. Age-specific properties of spontaneous mutations affecting mortality in *Drosophila melanogaster*. Genetics 148, 287–303.

Pletcher, S.D., Macdonald, S.J., Marguerie, R., Certa, U., Stearns, S.C., Goldstein, D.B., Partridge, L., 2002. Genome-wide transcript profiles in aging and calorically restricted *Drosophila melanogaster*. Curr. Biol. 12, 712–723.

Powell, S.J., Bale, J.S., 2005. Low temperature acclimated populations of the grain aphid *Sitobion avenae* retain ability to rapidly cold harden with enhanced fitness. J. Exp. Biol. 208, 2615–2620. doi:10.1242/jeb.01685

Promislow, D.E.L., Tatar, M., Khazaeli, A.A., Curtsinger, J.W., 1996. Age-specific patterns of genetic variance in *Drosophila melanogaster*. I. Mortality. Genetics 143, 839–848.

R Core Team, 2015. R: A language and environment for statistical computing. R Foundation for Statistical Computing.

Rajamohan, A., Sinclair, B.J., 2009. Hardening trumps acclimation in improving cold tolerance in *Drosophila melanogaster* larvae. Physiol. Entomol. 34, 217–223.

Reynolds, R.M., Temiyasathit, S., Reedy, M.M., Ruedi, E.A., Drnevich, J.M., Leips, J., Hughes, K.A., 2007. Age specificity of inbreeding load in *Drosophila melanogaster* and implications for the evolution of late-life mortality plateaus. Genetics 177, 587–595. doi:10.1534/genetics.106.070078

Rice, D.P., Townsend, J.P., 2012. Resampling QTL effects in the QTL sign test leads to incongruous sensitivity to variance in effect size. GenesGenomesGenetics 2, 905–911. doi:10.1534/g3.112.003228

Ricklefs, R.E., Finch, C.E., 1995. Aging: A Natural History, Scientific American Library Series.

Rieseberg, L.H., Widmer, A., Arntz, A.M., Burke, J.M., 2002. Directional selection is the primary cause of phenotypic diversification. Proc. Natl. Acad. Sci. 99, 12242–12245.

Rose, M.R., 1991. The Evolutionary Biology of Aging. Oxford University Press, Oxford.

Rose, M.R., 1984. Laboratory evolution of postponed senescence in *Drosophila melanogaster*. Evolution 38, 1004–1010.

Rose, M.R., Vu, L.N., Park, S.U., Graves, Jr., J.L., 1992. Selection on stress resistance increases longevity in *Drosophila melanogaster*. Exp. Gerontol. 27, 214–250.

Schnebel, E.M., Grossfield, J., 1988. Antagonistic pleiotropy: An interspecific *Drosophila* comparison. Evolution 42, 306. doi:10.2307/2409234

Schwarze, S.R., Weindruch, R., Aiken, J.M., 1998. Oxidative stress and aging reduce COX I RNA and cytochrome oxidase activity in *Drosophila*. Free Radic. Biol. Med. 25, 740–747.

Schwasinger-Schmidt, T.E., Kachman, S.D., Harshman, L.G., 2012. Evolution of starvation resistance in *Drosophila melanogaster:* Measurement of direct and correlated responses to artificial selection. J. Evol. Biol. 25, 378–387.

Sinclair, B.J., Roberts, S.P., 2005. Acclimation, shock and hardening in the cold. J. Therm. Biol. 30, 557–562. doi:10.1016/j.jtherbio.2005.07.002

Snoke, M.S., Promislow, D.E.L., 2003. Quantitative genetic tests of recent senescence theory: age-specific mortality and male fertility in *Drosophila melanogaster*. Heredity 91, 546–556. doi:10.1038/sj.hdy.6800353

Tatar, M., Promislow, D.E.L., Khazaeli, A.A., Curtsinger, J.W., 1996. Age-specific patterns of genetic variance in *Drosophila melanogaster*. II. Fecundity and its genetic covariance with age-specific mortality. Genetics 143, 849–858.

Vermeulen, C.J., Sørensen, P., Kirilova Gagalova, K., Loeschcke, V., 2013. Transcriptomic analysis of inbreeding depression in cold-sensitive *Drosophila melanogaster* shows upregulation of the immune response. J. Evol. Biol. 26, 1890–1902. doi:10.1111/jeb.12183

Wachter, K.W., Evans, S.N., Steinsaltz, D., 2013. The age-specific force of natural selection and biodemographic walls of death. Proc. Natl. Acad. Sci. 110, 10141–10146. doi:10.1073/pnas.1306656110

Wachter, K.W., Steinsaltz, D., Evans, S.N., 2014. Evolutionary shaping of demographic schedules. Proc. Natl. Acad. Sci. 111, 10846–10853. doi:10.1073/pnas.1400841111

Wickham, H., 2009. ggplot2: elegant graphics for data analysis. Springer New York.

Williams, G.C., 1957. Pleiotropy, natural selection, and the evolution of senescence. Evolution 11, 398. doi:10.2307/2406060

